# APLNR marks a cardiac progenitor derived with human induced pluripotent stem cells

**DOI:** 10.1101/2023.02.22.529606

**Authors:** Yin-Yu Lam, Chun-Ho Chan, Lin Geng, Nicodemus Wong, Wendy Keung, Yiu-Fai Cheung

## Abstract

Cardiomyocytes can be readily derived from human induced pluripotent stem cell (hiPSC) lines, yet its efficacy varies across different batches of the same and different hiPSC lines. To unravel the inconsistencies of in vitro cardiac differentiation, we utilized single cell transcriptomics on hiPSCs undergoing cardiac differentiation and identified cardiac and extra-cardiac lineages throughout differentiation. We further identified APLNR as a surface marker for in vitro cardiac progenitors and immunomagnetically isolated them. Differentiation of isolated in vitro APLNR^+^ cardiac progenitors derived from multiple hiPSC lines resulted in predominantly cardiomyocytes accompanied with cardiac mesenchyme. Transcriptomic analysis of differentiating in vitro APLNR^+^ cardiac progenitors revealed transient expression of cardiac progenitor markers before further commitment into cardiomyocyte and cardiac mesenchyme. Analysis of in vivo human and mouse embryo single cell transcriptomic datasets have identified APLNR expression in early cardiac progenitors of multiple lineages. This platform enables generation of in vitro cardiac progenitors from multiple hiPSC lines without genetic manipulation, which has potential applications in studying cardiac development, disease modelling and cardiac regeneration.

## Introduction

Cardiomyocytes can be readily generated from hiPSCs by temporal modulation of the Wnt pathway^1^. However, widespread applications for hiPSC derived cardiomyocytes, are hampered by unpredictable differentiation efficacy across multiple hiPSC lines^2^. Multiple studies have attempted demonstrated the presence of off-target endodermal and ectodermal cell lineages during in vitro cardiac differentiation of multiple hiPSC lines^3,4^. However, there has yet to be attempts to eradicate these cell populations to optimize cardiac differentiation in hiPSC lines with low differentiation efficacy. This is compounded by the lack of studies that compare cell populations between in vitro cardiac differentiation with in vivo cardiac development. Therefore, the potential of in vitro cardiac differentiation in furthering our understanding in vivo cardiac development and vice versa has yet to be unveiled.

Multiple surface markers have been proposed to isolate cardiovascular or cardiac progenitors and improve cardiac differentiation efficacy^5–8^. However, as the studies utilized only 1 to 2 human embryonic stem cell (hESC) or hiPSC lines, the possibility of variations and inconsistencies between different lines have yet to be accounted. This in turn limits widespread use of such markers for consistent generation of hESC or hiPSC derived cardiac lineages. Furthermore, as the isolated progenitors were only retrospectively characterized by their lineage descendants, it is not known which stage the isolated progenitors correspond to in vivo and if they represent a homogenous cell population.

In this study, we utilized single cell RNA sequencing (scRNA-seq) to dissect the transient cell populations across in vitro cardiac differentiation from different hiPSC lines and identify an in vitro cardiac progenitor population. We identified APLNR as a surface marker for in vitro cardiac progenitors and immunomagnetically isolated them. Isolated in vitro APLNR^+^ cardiac progenitors differentiate into predominantly cardiomyocytes accompanied with cardiac mesenchyme with similar efficacy in 3 hiPSC lines. Further transcriptomic analysis on differentiating in vitro APLNR^+^ cardiac progenitors consistently demonstrated transient expression of cardiac progenitor markers upon further commitment into cardiomyocyte and cardiac mesenchyme cell lineages in 3 hiPSC lines. We then analyzed published scRNA-seq datasets of a Carnegie Stage (CS) 7 human embryo and embryonic day (E) 7.75, E8.25 and E9.25 mouse hearts to identify the role of APLNR expressing cardiac progenitors in in vivo cardiac development^9,10^. Both datasets identified *APLNR*/*Aplnr* expression in human and mouse cardiac progenitors. Pseudotime analysis suggested *Aplnr* expressing cardiac progenitors differentiates into both multiple cardiac lineages at different time points. Our platform establishes a scalable and genetic manipulation free method of isolating in vitro cardiac progenitors from multiple hiPSC lines with consistent cardiac differentiation dynamics and efficacy, with potential applications in mechanistic study of early cardiac development, disease modelling and cardiac regeneration.

## Results

We first performed 16 batches of in vitro cardiac differentiation with an embryoid body based protocol on 3 hiPSC lines and calculated coefficient of variations of 37.7%, 16.0% and 41.0% in hiPSC 1, 2 and 3 respectively for cTnT^+^ cardiomyocytes at day 15 of cardiac differentiation^11^ (Figure 1A). This indicates significant variations in the efficacy of cardiac differentiation across different batches of the same hiPSC and different hiPSC lines. To dissect the variations in in vitro cardiac differentiation at a higher resolution, we then performed scRNA-seq on 23,511 cells from 2 hiPSC lines during day 2, 4, 5 and 9 of in vitro cardiac differentiation (Figure 1B). Clustering and UMAP analysis identified 10 cell populations, 7 cardiac and 3 extra-cardiac lineages present in both hiPSC lines. Moreover, we also identified a trajectory marking the transcriptomic changes from pluripotency to three cardiac lineage descendants: Cardiomyocytes, cardiac mesenchyme and endothelium (Figure 1C, Figure S1A). Analysis of differentially expressed genes (DEG) of each cell cluster demonstrated hiPSCs (*POU5F1, SOX2*) undergo mesodermal differentiation upon BMP and TGF-β stimulation via a transient mesendoderm (*MIXL1, NODAL*) population at day 2 before committing to a definitive mesoderm (*MESP1, CYP26A1*) population at day 4. This is followed by cardiac specification with Wnt antagonists to a cardiac progenitor (*HAND1, CFC1*) population at day 5 and further differentiation into cardiomyocyte (*TNNT2, NKX2-5*), cardiac mesenchyme (*COL3A1, COL1A2*) and endothelial (*ECSCR, CDH5*) populations at day 9 (Figure 1D, Figure S1B). This chronology is similar to those observed in in vivo mouse cardiac development, encompassing the biological processes of gastrulation, mesodermal differentiation, specification into cardiac lineages and differentiation into multiple cardiac cell populations^12^. This suggests cell populations present across in vitro cardiac differentiation are consistent between hiPSC lines and the transition cell identity across in vitro cardiac differentiation are similar to in vivo cardiac development.

**Figure 1.**
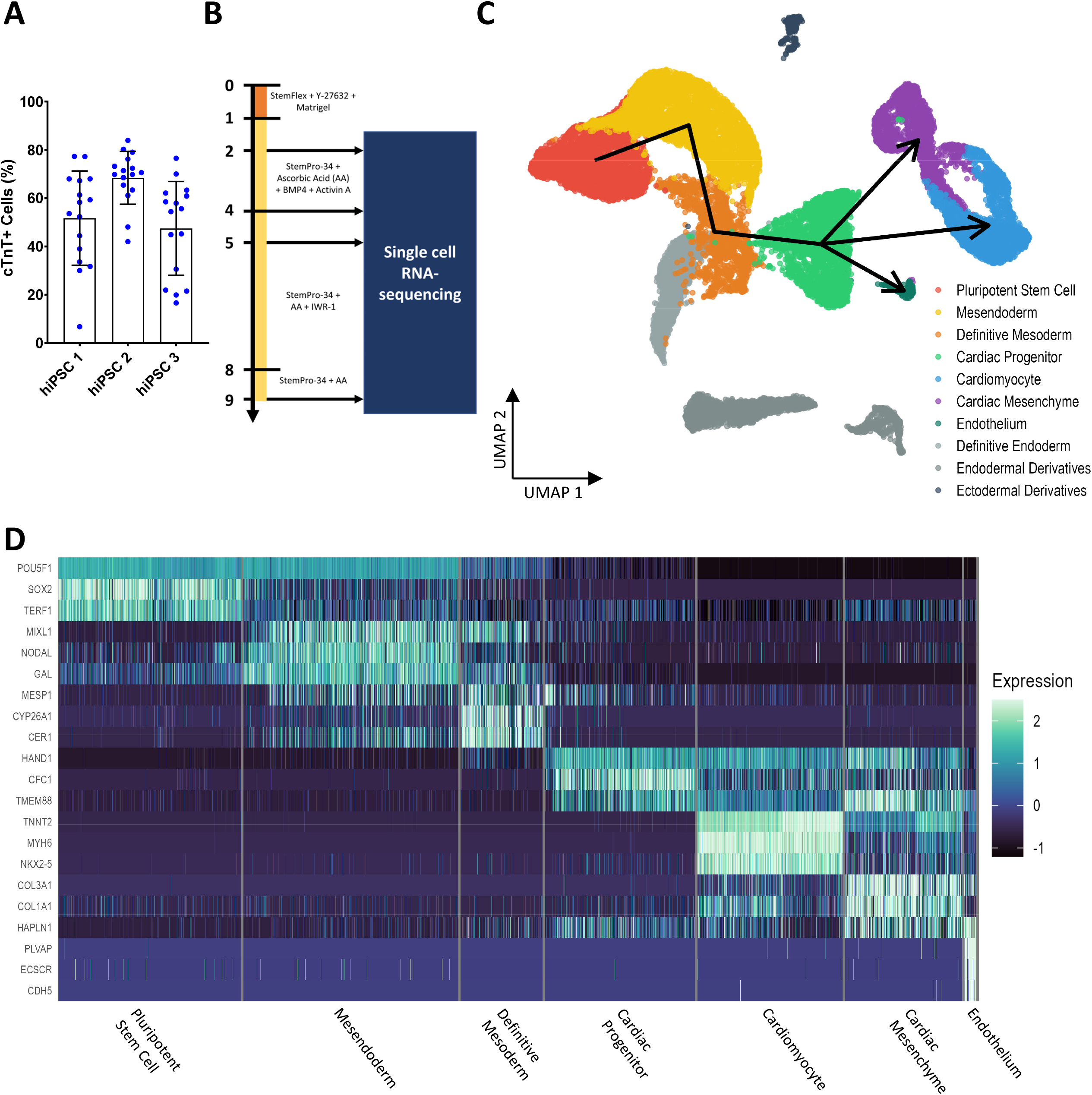
Transcriptomic changes across in vitro cardiac differentiation. A. Significant batch and line variations in cTnT^+^ cells at day 15 of in vitro cardiac differentiation. B. Experimental workflow to perform scRNA-seq sequentially on hiPSCs during different stages of in vitro cardiac differentiation from one differentiation each for hiPSC 1 and 2. C. UMAP representation of the scRNA-seq dataset with superimposed colours representing different cell clusters and a black arrow indicating the single cell trajectory of transcriptomic changes in in vitro cardiac differentiation. D. Single cell heatmap on the representative DEGs expressed in each cluster.

A mesenchymal or fibroblast-like cell of cardiac origin have been reported from single cell transcriptomics of in vivo embryonic mouse hearts and in vitro cardiac differentiation from multiple hiPSC and hESC lines^3,13,14^. However, the previous studies did not characterize this cell population in detail. DEG analysis between the cardiac mesenchyme and cardiomyocyte cell clusters showed higher expression of cardiac sarcomeric (*MYL3, TNNC1, MYH6, MYL7*) genes and cardiac transcription factors (*NKX2-5*) in cardiomyocytes and higher expression of extracellular matrix (*COL3A1, COL1A2, HAPLN1, LUM*) genes in cardiac mesenchyme (Figure 2A). Gene set enrichment analysis (GSEA) were consistent with DEGs in which cardiac muscle processes and development terms were enriched in the cardiomyocyte cluster while gene transcription and protein synthesis terms were enriched in the cardiac mesenchyme cluster (Figure 2B, 2C). This suggests cardiac mesenchyme is a distinct cell entity from cardiomyocytes and functions in the synthesis of extracellular proteins rather than muscle contraction. To further decipher the gene regulatory networks governing the transcriptomic differences between cardiomyocytes and cardiac mesenchyme, we then performed single cell regulon analysis on our scRNA-seq dataset at day 9 of cardiac differentiation containing the two cell clusters. UMAP and differential analysis of regulons revealed higher activity of cardiac transcription factors (*IRX4, MEF2C*) in cardiomyocytes and higher activity of transcription factors present in the cardiopharyngeal mesoderm (*LHX2*) and outflow tract (*PRDM6*) in cardiac mesenchyme^15,16^ (Figure 2D, 2E). This suggests cardiomyocyte and cardiac mesenchyme are two separate lineage descendants of in vitro cardiac progenitors with distinct transcriptomic identities.

**Figure 2.**
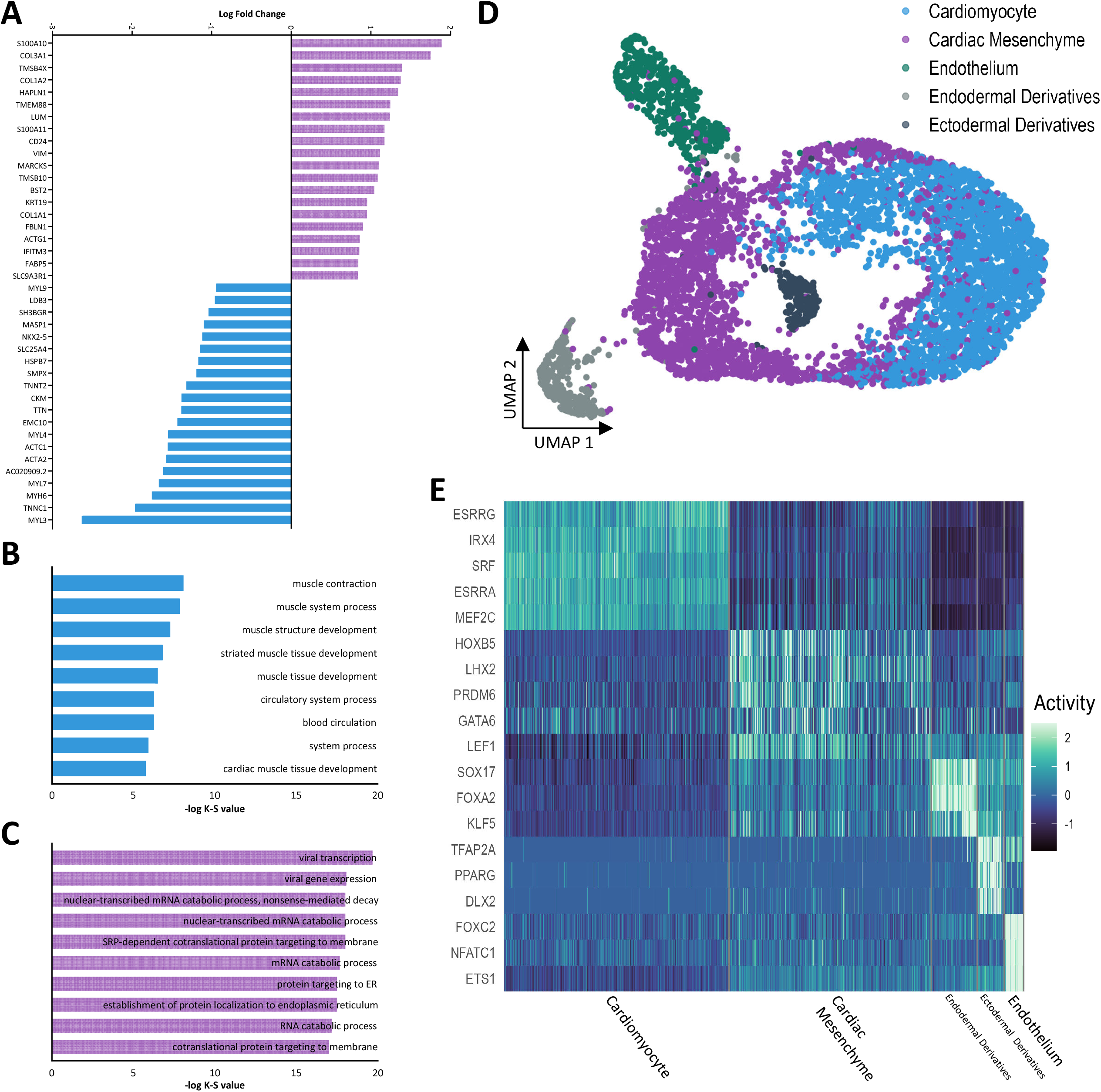
Cardiomyocyte and cardiac mesenchyme are distinct cell lineages derived from cardiac progenitors. A. Top 20 DEGs in cardiomyocytes and cardiac mesenchyme listed in descending order of logarithmic fold change. B. Enriched GO terms in the cardiomyocyte cluster listed in descending order of statistical significance. C. Enriched GO terms in the cardiac mesenchyme cluster listed in descending order of statistical significance. D. UMAP representation of regulons in the day 9 scRNA-seq dataset with superimposed colours representing clusters from figure 1B. E. Heatmap illustrating the top representative differential expressed regulons in day 9 clusters.

To further characterize the in vitro cardiac progenitor population, we performed DEG analysis on our scRNA-seq dataset at day 5 of cardiac differentiation consisting of the cardiac progenitor, definitive endoderm and pluripotent stem cell clusters (Figure 3A). *APLNR* was the 7th top DEG expressed in the cardiac progenitor cluster (Figure 3B). Its localization in the plasma membrane suggests it as a potential surface marker for positive isolation of in vitro cardiac progenitors. To evaluate *APLNR* with existing cardiac progenitor markers, we analyzed the expression of *APLNR* and known surface markers *KDR* and *PDGFRA*. All three markers were expressed in the cardiac progenitor cluster with *APLNR* having the highest expression, followed by *PDGFRA* and *KDR*. However, *KDR* and *PDGFRA* were also expressed in the pluripotent stem cell and definitive endoderm cell cluster respectively (Figure 3C). To validate our scRNA-seq analysis, we immunomagnetically isolated APLNR^+^ and APLNR^-^ cells at day 5 of cardiac differentiation and performed flow cytometry analysis of KDR and PDGFRA expression in the two cell fractions. Ubiquitous expression of KDR and PDGFRA are present in the APLNR^+^ cell fraction, while a population with weak KDR or PDGFRA is present in the APLNR^-^ cell fraction (Figure 3D, 3E). However, we also identified *APLNR* expression in mesodermal and cardiac lineages in day 4 and 9 of our scRNA-seq dataset, indicating the utility of APLNR in isolating in vitro cardiac progenitors is limited to day 5 of cardiac differentiation (Figure S2). Our findings are consistent with our scRNA-seq analysis and previous studies indicating KDR and PDGFRA marks extra-cardiac cell lineages^5,6^. Collectively, the findings indicate that APLNR is a surface marker for the in vitro cardiac progenitor population.

**Figure 3.**
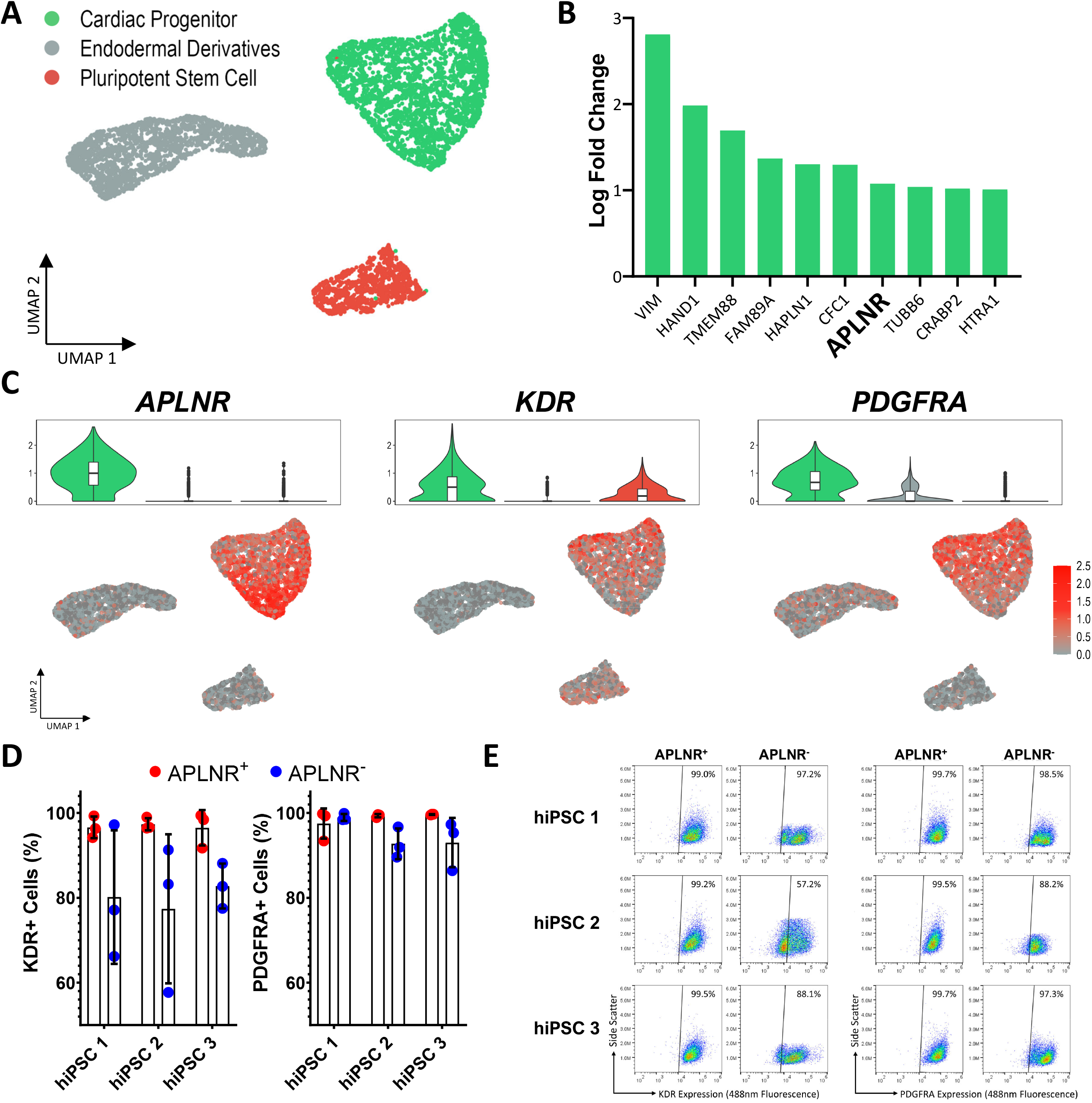
APLNR marks and immunomagnetically isolates cardiac progenitors. A. UMAP representation of the day 5 scRNA-seq dataset with superimposed colours representing different cell clusters. B. Top 10 DEGs in the cardiac progenitor cluster listed in descending order of logarithmic fold change. C. Top: Violin plot illustrating expression of cardiac progenitor surface markers in the dataset. Bottom: Single cell expression of the respective markers superimposed on the UMAP plot. D. Concurrent expression of KDR and PDGFRA in day 5 cells from the APLNR^+^ and APLNR^-^ cell fraction. Data represented as mean ± SD from three independent differentiations for each hiPSC line. E. Representative KDR (Left) and PDGFRA (Right) flow cytometry scatter dot-plots to illustrate two KDR^+^ and PDGFRA^+^ cell populations.

We next investigated the lineage descendants of in vitro APLNR^+^ cardiac progenitors by culturing them in unison with similar conditions compared to our cardiac differentiation protocol (Figure 4A). Flow cytometry analysis of cTnT on day 10 cells showed 75-80% cTnT^+^ cells in the APLNR^+^ cell fraction across all 3 hiPSC lines while variable cTnT expression was detected in the unsorted and APLNR^-^ cell fraction (Figure 4B, 4C). RT-qPCR analysis showed higher expression of cardiac progenitor, cardiomyocyte and cardiac mesenchyme genes in day 5 and day 10 APLNR^+^ cell fraction compared to unsorted control and APLNR^-^ cell fraction (Figure 4D, 4E). Immunofluorescence analysis also demonstrated higher and concentrated expression of cytoplasmic TNNT2 and extracellular COL3A1 in day 10 APLNR^+^ cell fraction (Figure 4F). Our findings collectively indicate that immunomagnetic positive sorting of APLNR^+^ cells at day 5 of cardiac differentiation isolates in vitro cardiac progenitors, which then differentiates into predominantly cardiomyocytes with accompanied cardiac mesenchyme cell populations in day 10 when cultured in isolation.

**Figure 4.**
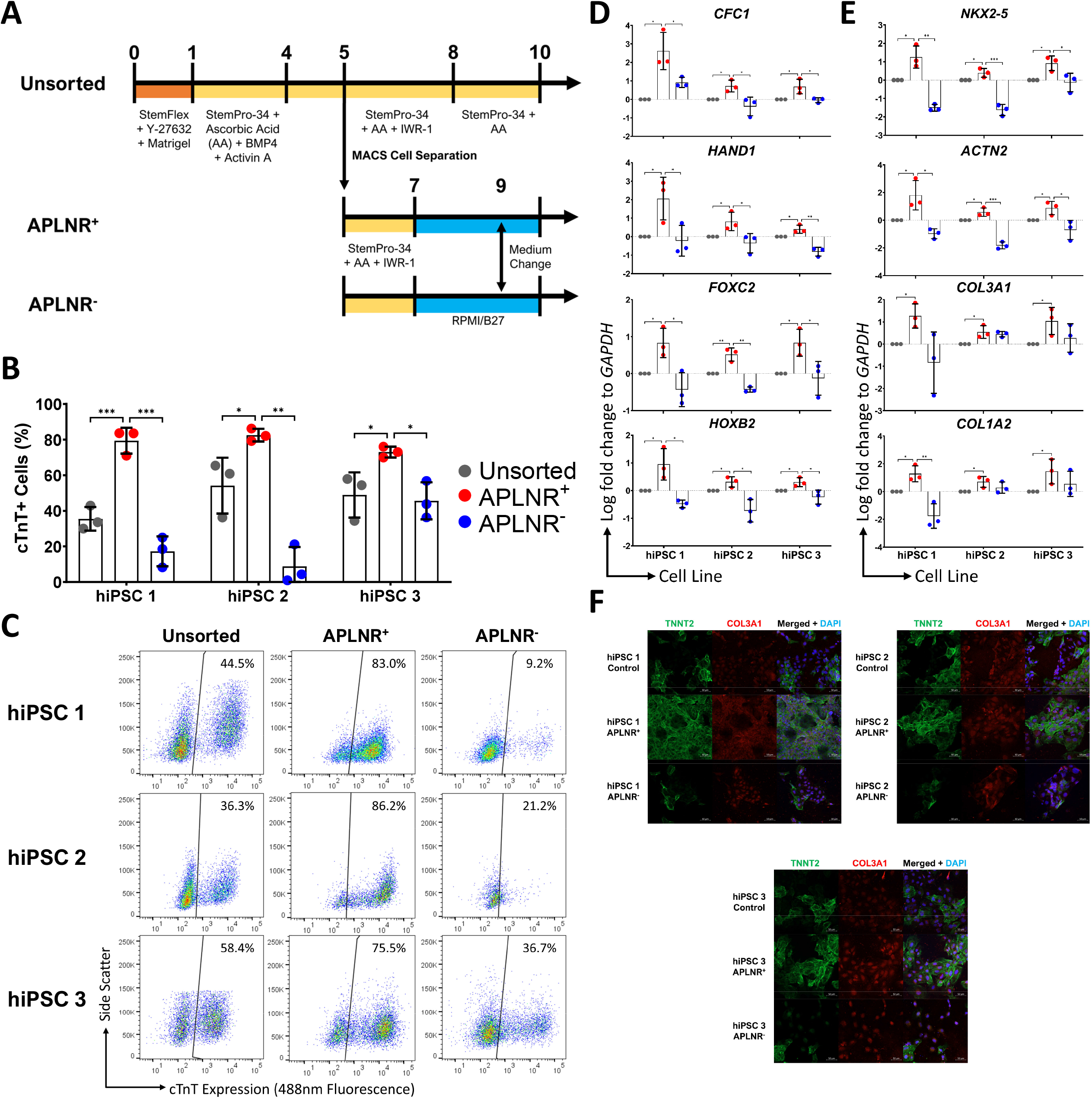
In vitro APLNR^+^ cardiac progenitors to differentiates into cardiomyocytes and cardiac mesenchyme. A. Experimental workflow to verify in vitro APLNR^+^ cardiac progenitors and its lineage descendants. B. Increased and consistent 75-80% cTnT^+^ in day 10 APLNR^+^ cell fraction. Data represented as mean ± SD from three independent differentiations for each hiPSC line. *P < 0.05, **P < 0.01, ***P < 0.001 C. Representative cTnT flow cytometry scatter dot-plots to illustrate figure 4B. D. Increased expression of cardiac progenitor markers in day 5 APLNR^+^ cell fraction. Data represented as mean ± SD from three independent differentiations for each hiPSC line. *P < 0.05, **P < 0.01, ***P < 0.001 E. Increased expression of cardiomyocyte and cardiac mesenchyme markers in day 10 APLNR^+^ cell fraction. Data represented as mean ± SD from three independent differentiations for each hiPSC line. *P < 0.05, **P < 0.01, ***P < 0.001 F. Immunocytochemistry showing Increased expression of TNNT2 and COL3A1 in day 10 APLNR^+^ cell fraction.

We next examined the differentiation trajectory of in vitro APLNR^+^ cardiac progenitors with RNA-seq on day 5, 6, 7 and 8 of cardiac differentiation (Figure 5A). Principal component analysis (PCA) identified differentiation time points as the major (65%) and inter-hiPSC line differences as the minor (15%) source of transcriptomic differences in our RNA-seq dataset. Furthermore, samples from all 3 hiPSC lines differentiate across a similar trajectory as defined temporally by PCA Dimension 1 (Figure 5B). This indicates in vitro APLNR^+^ cardiac progenitors and its lineage descendants until day 8 are transcriptomically consistent across multiple hiPSC lines. DEG analysis has identified downregulation of cardiac progenitor markers at day 6 accompanied with upregulation of cardiomyocyte and cardiac mesenchyme markers from day 6 to 8 (Figure 5C). GSEA has progressively enriched an increased number and statistical significance of cardiac muscle and development processes from days 6 to 8 (Figure 5D). This suggests rapid loss of progenitor identity with progressive transition to cardiomyocyte and cardiac mesenchymal cell identity upon isolated culture of in vitro APLNR^+^ cardiac progenitors. Temporal analysis of single gene expression identified progressive downregulation of cardiac progenitor markers (*CFC1, HAND1, FOXC2*) from day 5 onwards, followed by expression of cardiac progenitor (*FGF10, TBX5, NR2F2, MAB21L2, TBX5*) genes at day 6 and 7. Prominent expression of cardiomyocyte transcription factor (*MEF2C, NKX2-5, ANKRD1, SMYD1*), sarcomeric (*MYH6, MYH7*) and mesenchymal (*COL3A1, COL1A1, COL1A2, CD24*) genes were present from day 7 onwards. Furthermore, there were inconsistent upregulation of ventricular (*IRX4*) and epicardial (*TBX18*) markers with minimal to negligible expression of ventricular and pacemaker (*MYL2, SHOX2*) markers at day 8 (Figure 5E). Our findings collectively indicate isolated culture of in vitro APLNR^+^ cardiac progenitors derived from multiple hiPSC lines results in consistent transient expression of lineage specific cardiac progenitor markers before further differentiation into cardiac mesenchyme and cardiomyocytes.

**Figure 5.**
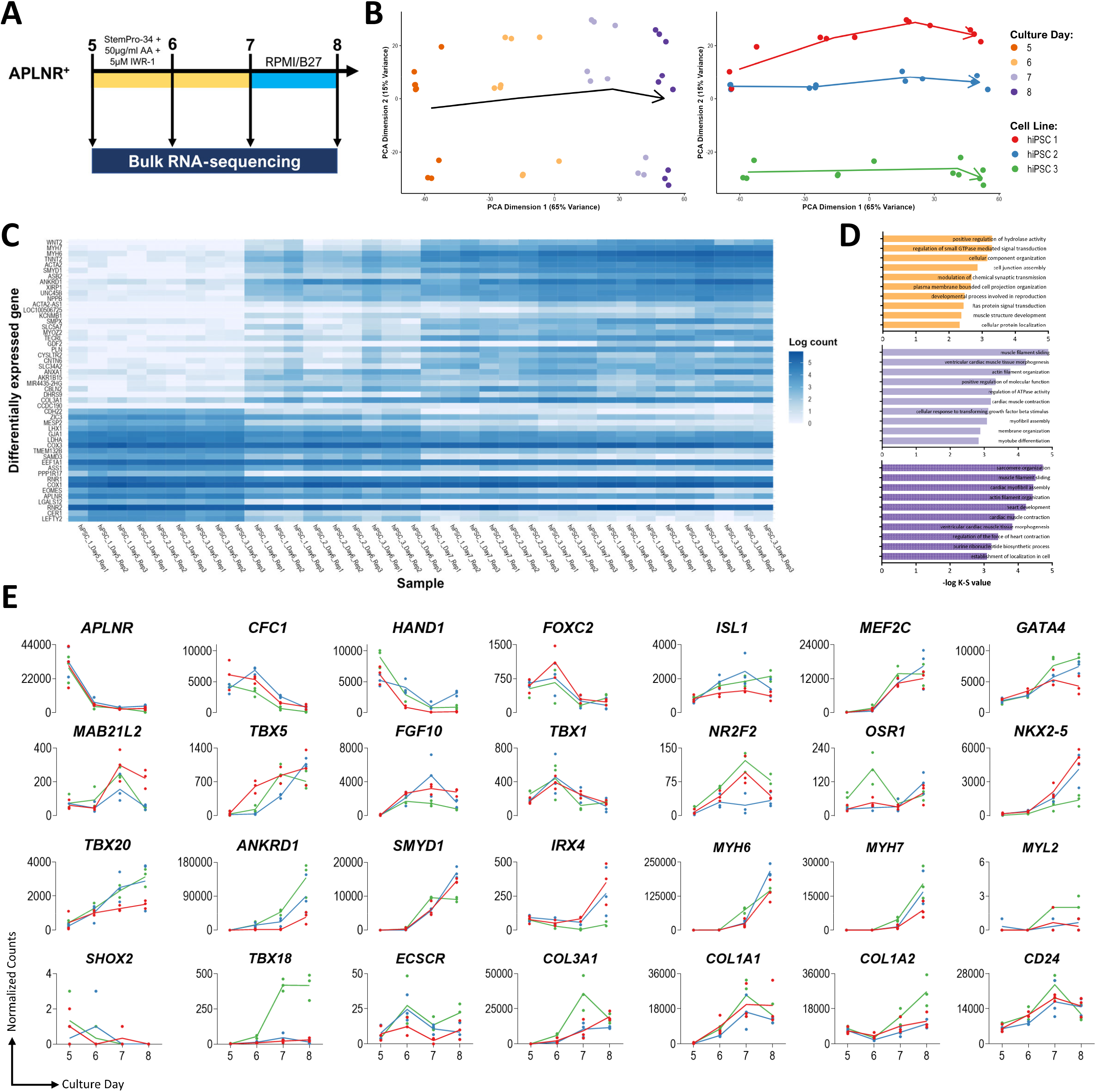
Transcriptomic changes of in vitro APLNR^+^ cardiac progenitors across time. A. Experimental workflow to examine the differentiation trajectory of in vitro APLNR^+^ cardiac progenitors with RNA-seq from three independent differentiations for each hiPSC line. B. PCA representation of the RNA-seq dataset with superimposed colours representing culture day (Left) and cell line (Right) and arrows indicating the trajectory of transcriptomic changes in differentiation of in vitro APLNR^+^ cardiac progenitors. C. Heatmap illustrating the top 50 DEGs in the dataset listed in descending order of fold change. D. Top 10 enriched GO terms in day 6 (Top), 7 (Middle) and 8 (Bottom) listed in descending order of statistical significance. E. Dot plot illustrating the individual expression of cardiac progenitor, lineage-specific progenitor, cardiomyocyte and their subtype, mesenchyme, epicardial and endothelial markers with superimposed colours representing hiPSC line of origin and respective coloured line connecting the arithmetic mean of expression over time for their respective hiPSC lines.

To validate the presence and explore the role of APLNR in in vivo cardiac development, we next analyzed a published scRNA-seq dataset consisting of 1,195 cells from a CS7 human embryo^9^. Clustering and UMAP analysis identified 10 cell populations consisting of pluripotent stem cells in the epiblast, derivatives and lineage descendants from all three germ layers (Figure 6A, 6B). *APLNR* expression was present in the primitive streak, definitive mesoderm, cardiac progenitor, mesenchyme and ciliated endoderm cluster, which is consistent with *APLNR* expression in in vitro cardiac progenitors identified from our scRNA-seq dataset (Figure 6C). To characterize the transcriptomic similarities between human in vivo and in vitro cardiac progenitors, we utilized scPred to classify individual cells from our in vitro cardiac differentiation scRNA-seq dataset with CS7 human embryo scRNA-seq dataset cell clusters based on transcriptomic similarity^17^. In vitro cardiac progenitors were classified predominantly to the human embryo cardiac progenitor cluster, with the rest in definitive mesoderm and primitive streak cluster. The in vitro cardiac mesenchyme cluster was partially classified into human embryo cardiac progenitor and mesenchyme cluster. In vitro cardiomyocytes were also partially classified into the human embryo cardiac progenitor cluster (Figure 6D, 6E). This suggests in vitro cardiac progenitors transcriptomically resemble CS7 human embryo cardiac progenitors and mesodermal progenitors entering the cardiac lineage, with in vitro cardiomyocytes and cardiac mesenchyme partially resembling human embryo cardiac progenitors. The findings collectively suggest *APLNR* is expressed in CS7 human embryo cardiac progenitors and are transcriptomically resemblant with predominantly in vitro cardiac progenitors with some cardiomyocytes and cardiac mesenchyme. This indicates the presence of a CS7 human embryo counterpart for in vitro APLNR^+^ cardiac progenitor and its cardiomyocyte and cardiac mesenchymal lineage descendants.

**Figure 6.**
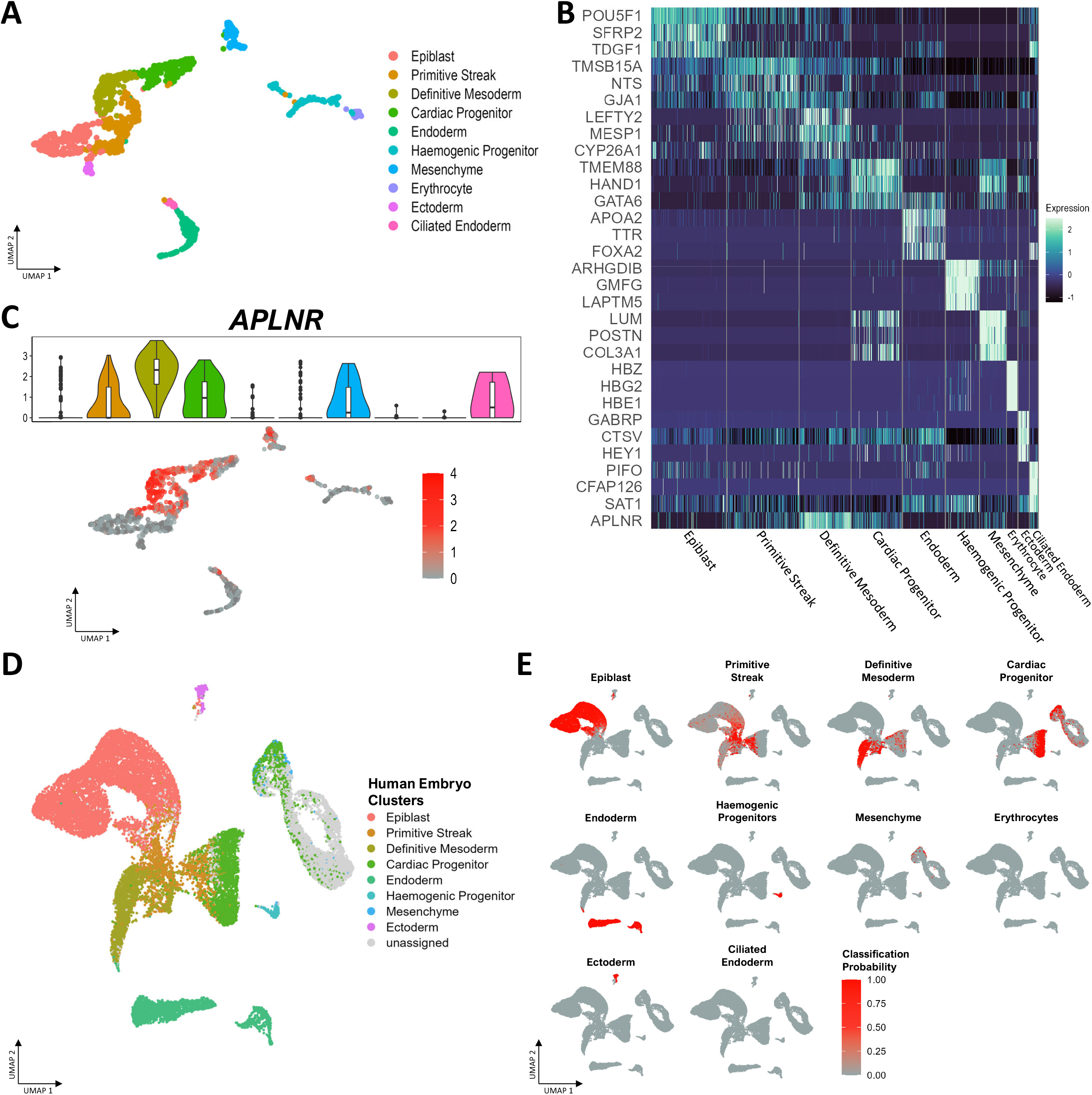
*APLNR* marks CS7 human cardiac progenitors. A. UMAP representation of the CS7 human embryo scRNA-seq dataset with superimposed colours representing different cell clusters. B. Single cell heatmap on the representative DEGs expressed in each cluster. C. Top: Violin plot illustrating expression of *APLNR* in the dataset. Bottom: Single cell expression of *APLNR* superimposed on the UMAP plot. D. UMAP representation of the in vitro cardiac differentiation scRNA-seq dataset with superimposed colours representing each cell and their most resemblant CS7 human embryo scRNA-seq cluster. E. UMAP representation of the in vitro cardiac differentiation scRNA-seq dataset with superimposed heatmap illustrating the transcriptomic similarity of individual cells in the in vitro cardiac differentiation scRNA-seq dataset to their respective CS7 human embryo scRNA-seq clusters.

To further examine the potential cardiac lineage descendants of APLNR^+^ cardiac progenitors in an in vivo setting, we then analyzed another published scRNA-seq dataset consisting of 16,233 cells from E7.75, E8.25 and E9.25 mouse hearts due to ethical limitations in obtaining a human embryo of corresponding age^10^. Traditional view of cardiogenesis starts with mesodermal specification into the cardiac lineage with the combination of high BMP and low Wnt activity conferred by embryonic signalling gradients^18,19^. This is followed by diversification into multiple cardiac progenitor populations, the juxta cardiac field (JCF) giving rise to the epicardium and first heart field (FHF) and the second heart field (SHF)^20,21^. We re-analyzed the dataset by first segregating the cells into JCF and SHF lineages based on known markers from both lineages, as the JCF population was not characterized when the dataset was published (Figure S3A, B). Clustering and UMAP analysis of the JCF lineage identified 9 cell clusters consisting of cardiac progenitors (Cluster 1, 2, 4), cardiomyocytes (Cluster 3, 5, 6, 7), epicardium (Cluster 8) and proepicardium (Cluster 9) lineages (Figure 7A). Co-expression of *Aplnr* with cardiac progenitor (*Cfc1*) and JCF (*Mab21l2*) markers was present in cluster 1 and 2 at E7.75, which were then downregulated with upregulation of FHF (*Tbx5*) and cardiomyocyte (*Nkx2-5, Myh6*) markers in cluster 3, 4 and 5 at E8.25, followed by expression of ventricular (*Myl2*), atrial (*Myh6*^+^, *Myh7*^-^, *Myl2*^-^), pacemaker (*Shox2, Vsnl1*) and epicardial (*Tbx18, Wt1*) markers in cluster 6, 7 and 8 at E9.25 (Figure 7B). Pseudotime analysis using cluster 1 or *Aplnr*^+^ cardiac progenitors at E7.75 as the root node similarly demonstrates the temporal changes of cardiac progenitor, cardiomyocyte and epicardial genes from E7.75 to E9.25 as described in Figure 7B (Figure S3C, D). This suggests *Aplnr* is expressed in JCF progenitors at E7.75 which can differentiate into FHF cardiomyocytes of atrial, ventricular, pacemaker subtypes and epicardium in E8.25 and E9.25. Clustering and UMAP analysis of the SHF lineage identified 11 cell clusters consisting of cardiac progenitors (Cluster 1, 2, 3, 4, 7), cardiomyocytes (Cluster 5, 6, 8, 9), cardiopharyngeal (Cluster 10) and mesenchymal (Cluster 11) lineages (Figure 7C). Co-expression of *Aplnr* with cardiac progenitor (*Cfc1, Isl1*) and Anterior heart field (AHF) (*Tbx1, Fgf10*) was present in cluster 1 and 2 at E7.75, while *Cfc1* and *Tmem88* expression were absent in cluster 3 at E8.25 and cluster 4 at E9.25. *Aplnr*^+^ cardiac progenitors at E7.75 and E8.25 downregulate progenitor markers and express cardiomyocyte (*Nkx2-5, Myh6*) markers in cluster 5 and ventricular (*Myl2*) markers in cluster 6 at E8.25 and E9.25. *Aplnr*^+^ cardiac progenitors at E9.25 bifurcates into expressing posterior SHF (*Tbx5*),and atrial (*Myh6*^+^, *Myh7*^-^, *Myl2*^-^) and pacemaker (*Shox2, Vsnl1*) cardiomyocytes in cluster 7 and 8 or to cardiomyocyte (*Nkx2-5, Myh6*) at cluster 9 and then to ventricular (*Myl2*) cardiomyocytes at cluster 6 (Figure 7D). Pseudotime analysis using cluster 2 and 4 or *Aplnr*^+^ cardiac progenitors at E7.75 and E9.25 as the root node similarly demonstrates the temporal changes of cardiac progenitor and cardiomyocyte genes across E7.75 and E9.25 as described from Figure 7D (Figure S3E, F). This suggests that *Aplnr* is expressed in SHF progenitors from E7.75 to E9.25, where the E7.75 and E8.25 progenitors differentiate into ventricular cardiomyocytes while the E9.25 progenitors differentiate into atrial, ventricular, pacemaker cardiomyocyte and epicardium. The findings collectively suggest that *Aplnr* is expressed in cardiac progenitors of the mouse heart at multiple time points with the potential to enter JCF at E7.75 and SHF lineages at E7.75 and E9.25 and subsequently into atrial, ventricular, pacemaker cardiomyocytes and epicardium.

**Figure 7.**
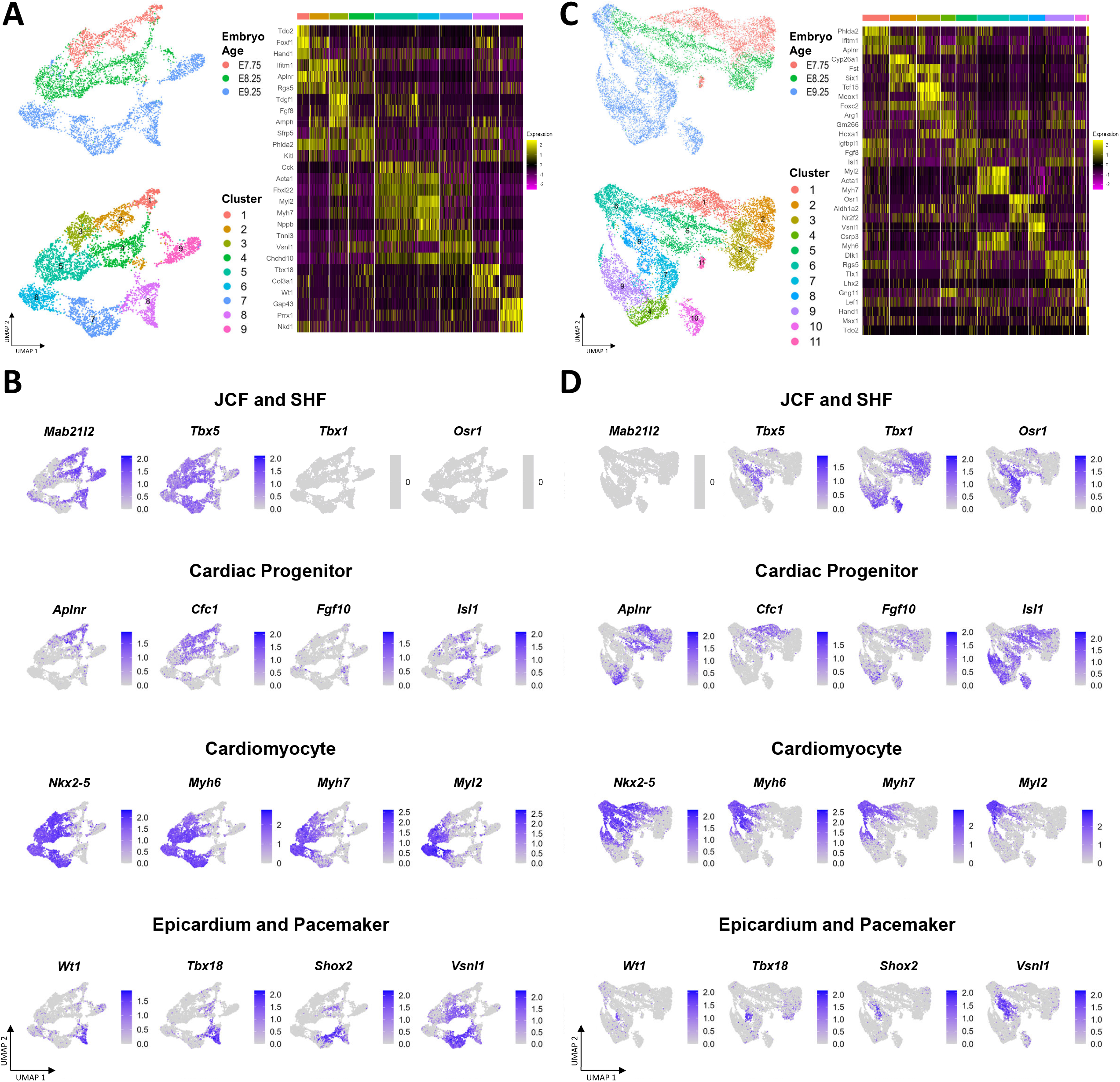
*Aplnr* marks in vivo mouse cardiac progenitors and differentiates into multiple cardiac lineages. A. Left: UMAP representation of the JCF lineage with superimposed colours representing embryo age (Top) and cell clusters (Bottom). Right: Single cell heatmap on the representative DEGs expressed in each cluster of the JCF lineage (Right). B. Single cell expression of cardiac progenitor, heart field lineages, cardiomyocyte and their subtypes and epicardial markers on the JCF lineage UMAP plot. C. Left: UMAP representation of the SHF lineage with superimposed colours representing embryo age (Top) and cell clusters (Bottom). Right: Single cell heatmap on the representative DEGs expressed in each cluster of the SHF lineage. D. Single cell expression of cardiac progenitor, heart field lineages, cardiomyocyte and their subtypes and epicardial markers on the SHF lineage UMAP plot.

## Discussion

Utilizing single cell transcriptomics on in vitro cardiac differentiation, we identified that cardiac and extra-cardiac cell populations are consistent across different hiPSC lines. We also identified APLNR as a surface marker for in vitro cardiac progenitors and immunomagnetically isolated them from multiple hiPSC lines, which then consistently differentiates into cardiomyocytes and cardiac mesenchyme with similar efficacy upon isolated culture. Transcriptomic analysis on differentiating in vitro APLNR^+^ cardiac progenitors identified temporally consistent transient expression of lineage specific markers similar to in vivo cardiac development before further differentiation into cardiomyocytes and cardiac mesenchyme in multiple hiPSC lines. Further analysis of a published CS7 human embryo scRNA-seq dataset similarly identified *APLNR* expression in human embryo cardiac progenitors and showed its transcriptomic congruence with in vitro cardiac progenitors. Analysis of a published E7.75, E8.25 and E9.25 mouse heart scRNA-seq dataset identified expression and potential of *Aplnr*^+^ cardiac progenitors to differentiate into JCF and SHF lineages, followed by further differentiation into epicardial and cardiomyocyte of atrial, ventricular and pacemaker subtypes.

Multiple studies have utilized single cell transcriptomics to characterize cell populations across in vitro cardiac differentiation. Previous studies have performed scRNA-seq on multiple time points across in vitro cardiac differentiation from two hiPSC lines and which identified endodermal lineages before and after Wnt inhibition and a non-contractile cardiomyocyte population with cardiomyocytes after day 15^3,4^. This is consistent with our findings in which we also identified endodermal lineages at day 4, 5 and 9 of our scRNA-seq dataset, suggesting the emergence of endodermal lineages from BMP or Wnt stimulation of hiPSCs is not an artefact, but due to partial or incomplete mesodermal specification with BMP or Wnt stimulation alone. Moreover, negligible expression of endodermal and ectodermal markers was found in our differentiating in vitro APLNR^+^ cardiac progenitors. In combination with its consistent differentiation efficacy across multiple hiPSC lines compared with our original embryoid body-based protocol, this suggests the presence of extra-cardiac lineages partially accounts for the variation in the efficacy of in vitro cardiac differentiation.

The presence of a cardiac lineage expressing extracellular matrix and mesenchymal genes is further verified by a large-scale study involving in vitro cardiac differentiation of 191 hiPSC lines. The study also reported variation in the proportion of mesenchymal lineages and cardiomyocytes across different hiPSC lines and suggested gene and X chromosome dosage skewing the cell fate determination of their common cardiac progenitor^14^. Further studies with scRNA-seq demonstrated the necessity of ISL1 expression during cardiac differentiation for generating a cardiac fibroblast-like cell population in one hESC line^22^. We also identified the presence of cardiac mesenchyme in all 3 hiPSC lines and generated it with similar efficacy from hiPSCs of both genders. This thereby does not support genetic factors affecting the balance of cardiomyocytes and cardiac mesenchyme. We further demonstrated with that cardiac mesenchyme and cardiomyocyte are descendants from APLNR^+^ cardiac progenitors with expression of their respective markers starting at day 7. Moreover, as APLNR expression precedes *ISL1* and other lineage specific cardiac progenitor genes in our RNA-seq dataset, this suggests cardiac mesenchyme is a distinct lineage derived from cardiac progenitors rather than a cardiomyocyte subtype.

The two-step approach of BMP or Wnt stimulation followed by Wnt inhibition is the backbone of in vitro cardiac differentiation for the past decade. Moreover, variations in applying the two-step approach to hiPSCs or hESCs were developed in the same time frame. The first is utilization of a two-dimensional monolayer versus three-dimensional embryoid body platform. Suspension culture of embryoid bodies is more resemblant to the developing embryo and amenable to scalability. However, difficulty in achieving consistent embryoid body size and led to the development of monolayer platforms at the expense of scalability^23^. The second is utilization of recombinant proteins or small molecules or combinations of the two to implement the two-step approach. Mesodermal specification have been achieved in different protocols by BMP or Wnt stimulation alone or a combination of the two, cardiac specification is further achieved with Wnt inhibition^1,5,24,25^. The transcriptomic similarities between our in vitro APLNR^+^ progenitors with CS7 human cardiac progenitors supports that embryoid bodies mimic embryonic development. Previous studies demonstrated the transcriptomic progression across in vitro cardiac differentiation are different between embryoid body and monolayer-based protocols^26^. Therefore, further studies will be required to validate the temporal profile and utility of in vitro APLNR^+^ progenitors in monolayer-based protocols and different cardiac induction methods.

Identification of surface markers for mesodermal and cardiac progenitor populations have been reported as a method to remove side lineages and improve cardiac differentiation efficacy. Isolation and culture of KDR^+^ c-KIT^-^ cardiovascular progenitors in two hESC lines results in a mix of 50% cardiomyocytes, 45% smooth muscle and 5% endothelium^5^. Increased efficacy was achieved via isolation of dual KDR^+^ PDGFRA^+^ cardiac progenitors, resulting in 80% cardiomyocytes in one hESC line. Moreover, the proportion of dual KDR^+^ PDGFRA^+^ or dual GFRA2^+^ PDGFRA^+^ cells was found to be predictive of cardiac differentiation efficacy^6,7^. Our day 9 scRNA-seq dataset showed similar proportions of cardiomyocytes, cardiac mesenchyme and endothelium, suggesting in vitro APLNR^+^ cardiac progenitors belong to the same lineage as KDR^+^ c-KIT^-^ cardiovascular progenitors. Furthermore, as KDR and PDGFRA are ubiquitously expressed in in vitro APLNR^+^ cardiac progenitors, the predictive nature of the dual markers can be due to direct or indirect measurement of the proportion of APLNR^+^ cardiac progenitors. This collectively suggests APLNR is comparable to dual KDR PDGFRA for isolating in vitro cardiac progenitors, with benefits of using one rather than two markers and validation of similar efficacies in multiple hiPSC lines. Other markers such as CXCR4, ITGA3 and CD1D have also been reported to enrich or isolate for SHF, AHF and pSHF lineage cardiac progenitors respectively in 1 hiPSC or hESC line^8,27^. As our in vitro APLNR^+^ cardiac progenitors transiently express multiple lineage-specific markers upon isolated culture, we are unable to deduce the proportion of lineage-specific cardiac progenitors. Hence, further studies on cell fate determination of in vitro APLNR^+^ cardiac progenitors will be required to make a comparison with existing lineage-specific markers.

All the aforementioned cardiac progenitor populations were all isolated via flow cytometry, as opposed to magnetic cell separation with in vitro APLNR^+^ cardiac progenitors. Magnetic cell separation offers a shorter processing time, cheaper setup with ease of scalability compared to than flow cytometry, which potentially makes in vitro APLNR^+^ cardiac progenitors a more practical choice for future applications. Furthermore, all the studies were performed on one or two hESC or hiPSC lines and hence the possibility of variations between different lines cannot be excluded. This point was alluded when >65% of dual KDR^+^ PDGFRA^+^ cardiac progenitors generated 82%, 60% and 37% cardiomyocytes in hES2, H1 hESCs and one hiPSC line respectively under optimized cardiac differentiation settings^6^. We have tested our in vitro APLNR^+^ progenitors on 3 hiPSC lines and demonstrated minimal inter-line differences on the cellular and transcriptomic level. This suggests the temporal expression of APLNR and cell identity of APLNR^+^ cells at day 5 of cardiac differentiation is consistent across multiple hiPSC lines.

In this study we uncovered the cell lineages across in vitro cardiac differentiation of hiPSCs with scRNA-seq. We also identified APLNR as a surface marker for a cardiac progenitor population and isolated it immunomagnetically. Transcriptomic analysis of differentiating in vitro APLNR^+^ cardiac progenitors derived from multiple hiPSC lines showed consistent differentiation via transient lineage-specific progenitors before differentiating into terminal cardiac lineages of similar efficacy. Further analysis of published human and mouse heart scRNA-seq datasets similarly identified *APLNR* expression in early cardiac progenitors from multiple lineages and time points. We have established a scalable platform for isolating in vitro cardiac progenitors from multiple hiPSC lines with minimal inter-line differences, which can be applied for disease modelling, studying early cardiac development and cardiac regeneration therapies.

## Supporting information

Key Resources Table

## Acknowledgements

We thank the Genomics & Bioinformatics Cores for providing assistance on scRNA-seq and RNA-seq and Imaging and Flow Cytometry Core of the Centre for PanorOmic Sciences at the Li Ka Shing Faculty of Medicine, The University of Hong Kong for flow cytometry and imaging support.

## Author contributions

Y.Y.L., C.H.C., W.K. and Y.F.C. conceived the study. G.L. and N.W. performed hiPSC culture and validation, G.L., Y.Y.L. and C.H.C. performed directed cardiac differentiation and flow cytometry validation. Y.Y.L. and C.H.C. performed immunomagnetic cell isolation and prepared cells for scRNA-seq. Y.Y.L. performed immunofluorescence imaging, prepared cells and performed computational analysis for RNA-seq and RT-qPCR datasets. Y.Y.L. and C.H.C. prepared cells and performed computational analysis for scRNA-seq datasets. Y.Y.L., W.K. and Y.F.C. wrote the manuscript.

## STAR Methods

### Resource Availability

#### Lead contact

Further information and requests for resources and reagents should be directed to and will be fulfilled by the lead contact, Professor Yiu-Fai Chueng.

#### Materials availability

This study did not generate new unique reagents.

#### Data and code availability

scRNA-seq and RNA-seq sequencing data have been deposited in GEO under the accession code GSE174213 and GSE174335 respectively.

### Experimental model and subject details

#### Generation of hiPSCs

Three healthy subjects (Two female, one male) were recruited with whole blood drawn. Peripheral blood mononuclear cells were then isolated from whole blood with SepMate™ (STEMCELL Technologies) and Lymphoprep™ (STEMCELL Technologies). Peripheral CD34^+^ haematopoietic progenitors were then isolated with the Human CD34 MicroBead Kit (Miltenyi Biotec) according to manufacturer’s protocol, which were then cultured in StemSpan™ H3000 (STEMCELL Technologies) with CC100 (STEMCELL Technologies) for 3 days. Episomal reprogramming was the reprogrammed by nucleofection of pCXLE-hOCT3/4-shp53, pCXLE-hSK and pCXLE-hUL with the Human CD34 Cell Nucleofector™ Kit (Lonza) according to manufacturer’s protocol^28^. Nucleofected putative hiPSCs were then maintained in Geltrex™ (Gibco) coated 6-well plates in StemFlex™ (SF; Gibco) medium. hiPSC purification was then performed with human Anti-TRA-1-60 Microbeads (Miltenyi Biotec) after two weeks in culture.

hiPSCs were validated by immunofluorescence (IF) staining of pluripotency markers. hiPSCs were fixed with 4% paraformaldehyde (PFA) in phosphate-buffered saline (PBS)(Affymetrix) for 15 minutes in room temperature, followed by permeabilization with 0.1% Triton X-100 (Sigma-Aldrich) in PBS for 10 minutes in room temperature. Fixed hiPSCs were then immunostained with pluripotency markers: anti-OCT3/4 (Santa Cruz Biotechnology; SC-5279, 1:200), anti-SOX2 (Santa Cruz Biotechnology; SC-17320, 1:100), anti-SSEA4 (STEMCELL Technologies; MC-813-70, 1:200) or anti-TRA-1-81 (Cell Signalling Technology; 4745S, 1:100) at 4°C overnight, followed with Alexa Fluor® (AF)-488 conjugated donkey anti-goat IgG (Invitrogen; A-11055, 1:200) or AF-488 conjugated donkey anti-mouse IgG (Invitrogen; A-21202, 1:200) secondary antibodies at 4°C for 1 hour and DAPI (Invitrogen) for 5 minutes in room temperature. Stained cells were mounted with ProLong™ Gold Antifade Mountant (Invitrogen). Imaging was then performed with LSM 700 (Carl Zeiss). Validation images are available in our previous publication^29^.

### Method details

#### Directed cardiac differentiation for hiPSCs

hiPSCs were grown to 80% confluency and dissociated into single cells with StemPro™ Accutase™ (Gibco), which were transferred into ultra-low attachment six-well plates (Corning) and cultured in SF with 40µg/ml Matrigel™ (Corning), 1ng/ml BMP4 (Gibco) and 10µM Y-27632 (BioGems) under a hypoxic 5% O_2_ environment and defined as day 0. Culture medium was switched to StemPro™-34 SFM (SP34; Gibco) with 50µg/ml ascorbic acid (AA; Sigma-Aldrich), 2mM GlutaMAX™ (Gibco), 10ng/ml BMP4 and 10ng/ml Activin A (Gibco) in day 1. BMP4 and Activin A were replaced with 5µM IWR-1 (STEMCELL Technologies) in day 4. IWR-1 was then removed, cells were transferred into a normoxic environment and maintained in SP34 with GlutaMAX™ and AA replaced twice per week at day 8 onwards. Day 15 cells were dissociated with 0.025% Trypsin-EDTA (Gibco) for flow cytometry analysis.

#### Immunomagnetic isolation of APLNR^+^ progenitors

Day 5 cells were harvested and dissociated into single cells with TrypLE™ Express (Gibco), which were then immunostained with anti-hAPJ (R&D MAB8561; 1:200) in Hanks’ Balanced Salt Solution (HBSS; Gibco) + 0.5% Bovine serum albumin (BSA; Sigma-Aldrich) at room temperature for 30 minutes followed with goat anti-mouse IgG microbeads (Miltenyi Biotec) at room temperature for 15 minutes. Positive selection was then performed with MS columns (Miltenyi Biotec) and OctoMACS™ Separator (Miltenyi Biotec). APLNR^+^ cells were then plated on Matrigel™ coated 24-well plates at a density of 3.5 - 4 × 10^5^ cells/well and cultured with day 4 cardiac differentiation medium for two days followed by RPMI 1640 with GlutaMAX™ (Gibco) and 2% B27 supplement (Gibco) replaced every other day. Day 10 cells were dissociated with 0.025% Trypsin-EDTA for subsequent analysis.

#### RT-qPCR

Total RNA was isolated with TRIzol™ LS Reagent (Invitrogen) according to manufacturer’s protocol, which is then quantified with NanoDrop™ 2000c (Thermo Fisher Scientific). cDNA synthesis was performed with 1µg of total RNA using QuantiTect™ Reverse Transcription Kit (Qiagen) according to manufacturer’s protocol. cDNA quantification was performed in 96-well PCR plates with 20µl reaction volume using LightCycler® 480 (Roche). Each reaction consists of 10ng of cDNA template, 400nM of forward and reverse primers and iTaq™ Universal SYBR Green Supermix (Bio-Rad). The plates were incubated at 95°C for 30 seconds, followed by 40 cycles of 95°C for 5 seconds and 60°C for 30 seconds. Relative gene expression was calculated with the 2^-^ΔΔ^Ct^ method normalized to GAPDH expression. Primer sequences are available at Supplementary Table 1.

#### RNA-seq

Total RNA was isolated with TRIzol™ LS Reagent (Invitrogen) according to manufacturer’s protocol, which is then quantified with NanoDrop™ 2000c and integrity checked with 2100 BioAnalyzer (Agilent) using the RNA 6000 Nano Kit (Agilent). Poly-A mRNA-seq was then performed with NovaSeq 6000 System (Illumina). Raw reads were then aligned against the GRCh38_p13 genome with Rsubread^30^. Subsequent analysis with performed DESeq2^31^. PCA analysis was performed with the plotPCA function and DEGs were computed by comparing day 6, 7 or 8 with the day 5 dataset. Genes with > 2 logarithmic fold change and an adjusted P-value < 0.01 was selected for GSEA analysis using topGO.

#### scRNA-seq

For the in vitro cardiac differentiation dataset, day 2, 4, 5 and 9 cells undergoing cardiac differentiation were dissociated into single cells with TrypLE™ Express followed by cell encapsulation and library preparation with Chromium Single Cell 3’ Reagent Kit v3 (10X Genomics). Library sequencing was performed with the NovaSeq 6000 System. Raw reads were then aligned and initially filtered with Cell Ranger (10X Genomics). Analysis was performed with Seurat 3.0^32^. Cell filtering was performed by eliminating cells with low number of RNA transcripts and high mitochondrial gene percentage followed by logarithmic normalization. Top 2000 most variable genes were chosen to compute the PCA. The number of PCAs used for k-means clustering analysis and Uniform Manifold Approximation and Projection (UMAP) visualization was determined with the Scree test. DEG analysis was calculated with the FindAllMarkers function and only genes expressed in at least 25% of the cell cluster, > 0.5 logarithmic fold change and an adjusted P-value < 0.05 were included. Trajectory analysis was performed with connecting cluster centroids based on biological significance. Regulon analysis was performed with pySCENIC^33^. Genes that were expressed in more than 1% of the cells were selected, regulons were then ranked with the GRCh38 RefSeq r80 database consisting of 10kb upstream and downstream and 500bp upstream and 100bp downstream of the transcriptional start site. Results were then imported into Seurat 3.0, PCA, clustering and UMAP analysis was performed as above and differential regulons were calculated with the same manner as DEGs.

For the CS7 human embryo dataset, we obtained the raw counts from dataset was performed from ArrayExpress under the accession code E-MTAB-9388. Adapter trimming on raw reads was done with cutadapt, followed by alignment with the GRCh38_p13 genome with STARsolo^34,35^. Subsequent cell filtering, normalization, PCA, clustering, UMAP and DEG analysis was performed with Seurat 3.0 using the same methods as the in vitro cardiac differentiation dataset. Classification with the CS7 human embryo clusters was performed with scPred using the Mixed Determinant Analysis model^17^.

For the E7.75, E8.25 and E9.25 mouse heart dataset, the expression matrix was directly obtained from GEO under the accession code GSE126128. Initial cell filtering, normalization, top 800 variable genes used to compute PCA and UMAP analysis was performed with Seurat 3.0 using the same methods as the in vitro cardiac differentiation dataset. Cells were determined to be of Juxta cardiac field (JCF) lineage if they express *Mab21l2* or *Tbx5* and without *Tbx1* and *Osr1* expression, those are of Second heart field (SHF) lineage if they are excluded from the JCF lineage and without *Mab21l2* expression. Subsequently cell cycle genes were regressed, the top 1000 genes were to used compute PCA, clustering, UMAP and DEG analysis with Seurat 3.0 using the same methods as the in vitro cardiac differentiation dataset. Pseudotime analysis was performed with Monocle 3 using the cluster containing *Aplnr*^+^ cardiac progenitors as the root node^36^.

#### Immunofluorescence imaging

Dissociated cells were cultured on 12mm glass cover slips (Marienfield), which were subsequently fixed with 4% PFA for 15 minutes at room temperature, followed by permeabilization with 0.1% Triton X-100 in PBS for 10 minutes at room temperature. Fixed cells were then stained with anti-cTnT (Invitrogen; MA5-17192, 1:400), anti-COL3A1 (Novus Biologicals; NB600-594, 1:200) at 4°C overnight, followed with AF-488 conjugated goat anti-mouse IgG (Invitrogen; A-10680, 1:1000) and AF-555 conjugated goat anti-rabbit IgG (Invitrogen; A-21428, 1:1000) secondary antibodies at room temperature for 1 hour and 300nM DAPI (Invitrogen; D1306) for 5 minutes in room temperature. Stained cells were mounted with ProLong™ Gold Antifade Mountant. Imaging was then performed with LSM 800 (Carl Zeiss).

#### Flow cytometry

Dissociated cells were fixed and permeabilized with BD Cytofix/Cytoperm™ (BD Biosciences) according to manufacturer’s protocol. Fixed cells were then stained with anti-cTnT (1:400), anti-KDR (Cell Signalling Technology; 9698S; 1:200), anti-PDGFRA (Cell Signalling Technology; 3164S; 1:200) antibodies at 4°C overnight followed by FITC conjugated rat anti-mouse IgG_1_ antibody (BioLegend; 406605, 1:50) or Alexa Fluor 488 conjugated goat polyclonal anti-rabbit IgG (H+L) (Abcam; ab150077; 1:1000) at 4°C for 1 hour. Stained cells were counted with FACSCanto™ II (BD Biosciences). Analysis was performed with FlowJo (FlowJo LLC).

### Quantification and statistical analysis

Statistical analysis was performed with GraphPad Prism 9.0. All descriptive statistics are reported as mean ± SD. Statistical significance was evaluated with two-tailed t-test and p value < 0.05 was considered as significant unless stated otherwise.

