## Supplementary Materials for "APLNR marks a cardiac progenitor derived with human induced pluripotent stem cells"

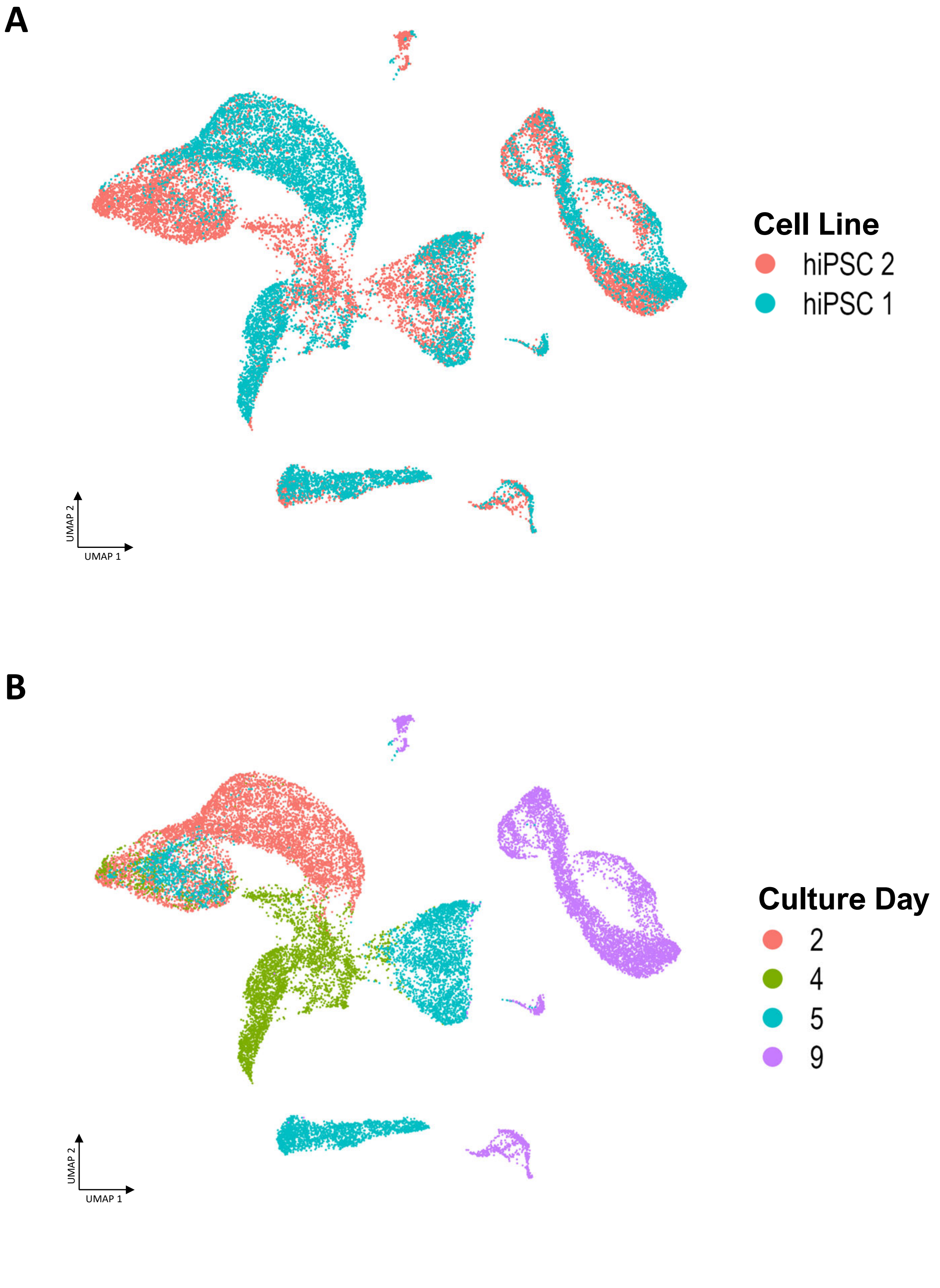


**Supplementary Figure 1. UMAP representation of the in vitro cardiac differentiation scRNA-seq dataset with different variables**

A. Superimposed colours representing different hiPSC lines.

B. Superimposed colours representing culture day when sample was sequenced.

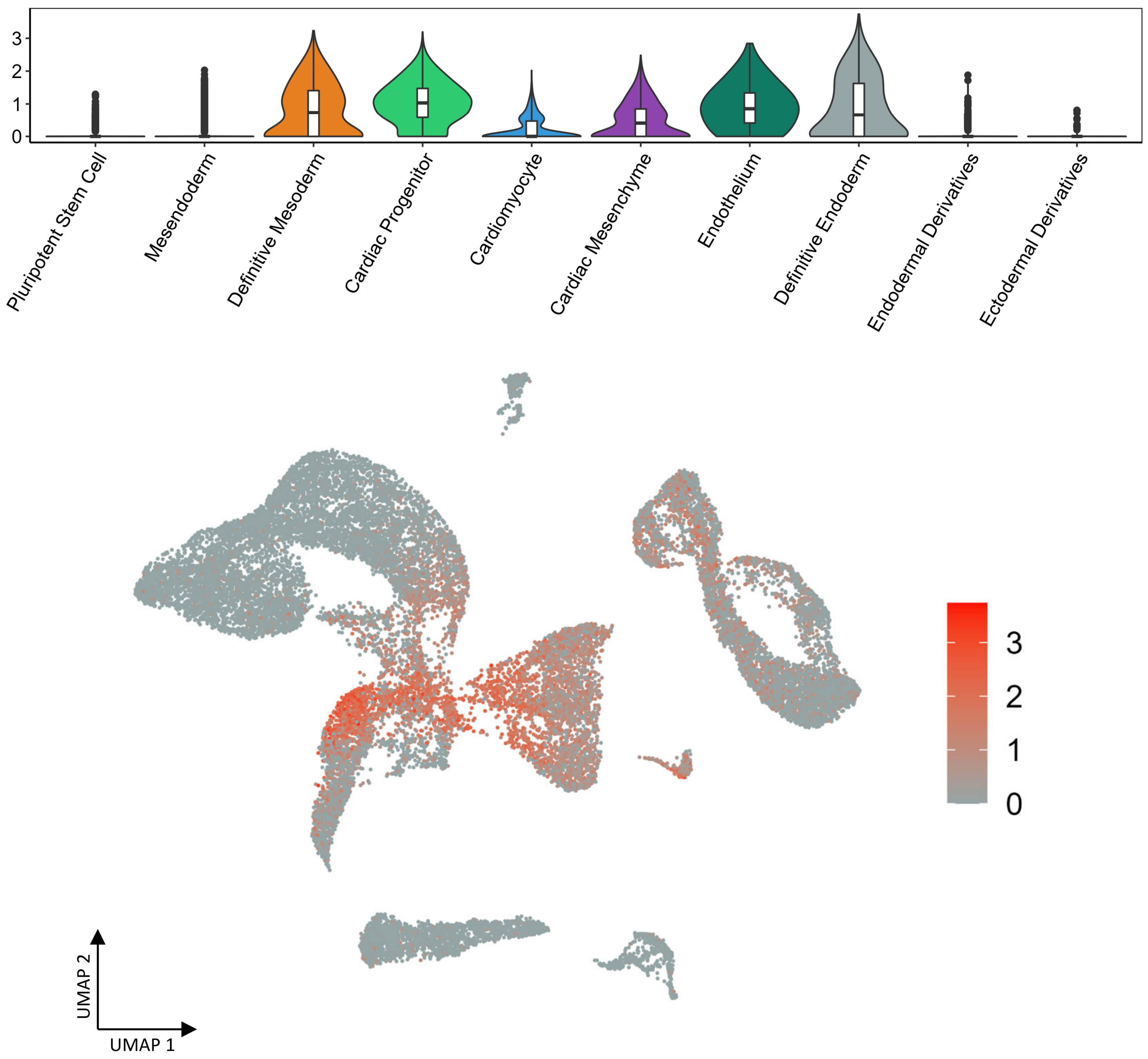


**Supplementary Figure 2. *APLNR* expression in the in vitro cardiac differentiation scRNA-seq dataset**

Top: Violin plot illustrating expression of *APLNR* in the dataset. Bottom: Single cell expression of *APLNR* superimposed on the UMAP plot.

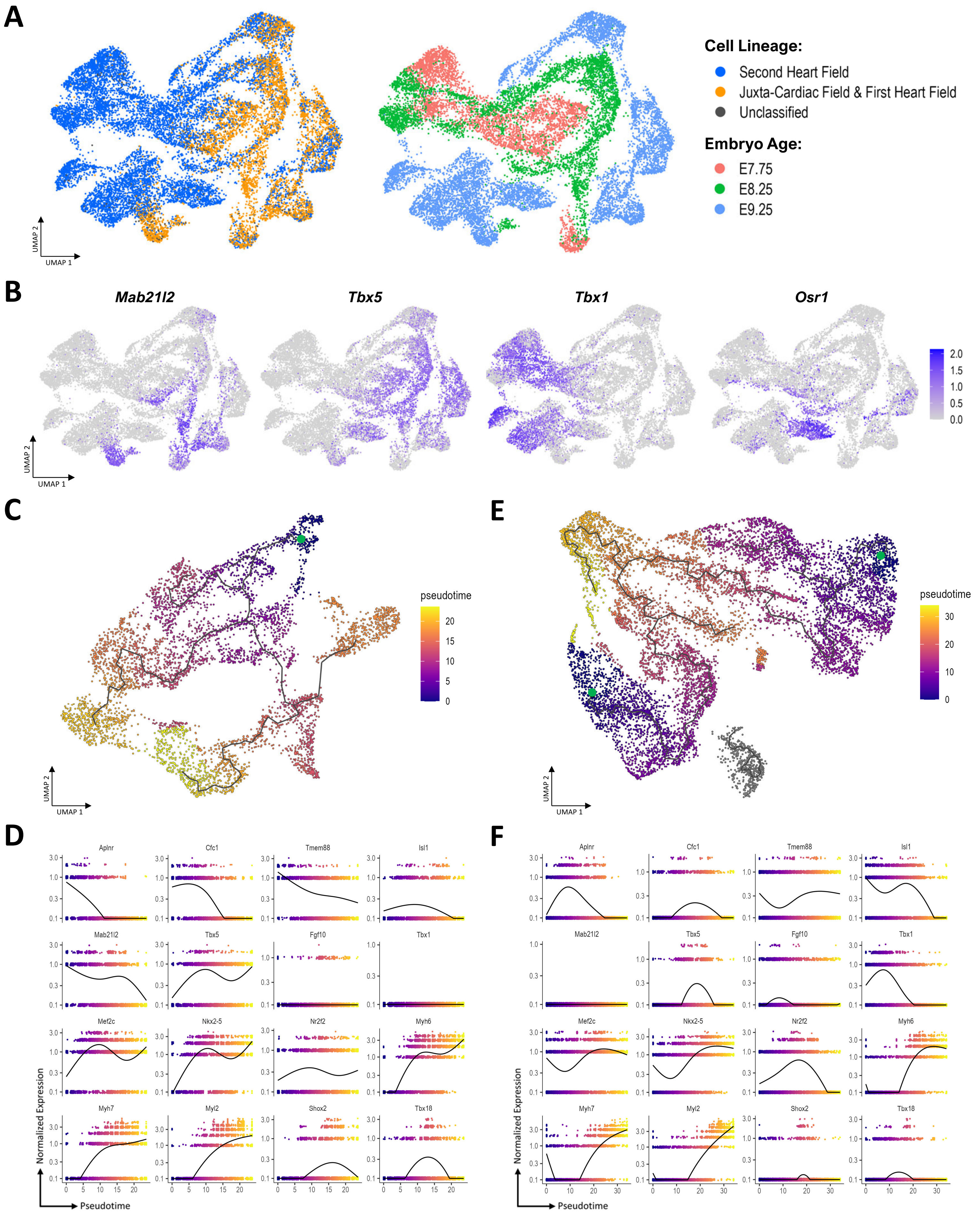
A
Cell Lineage:
Second Heart Field
Juxta-Cardiac Field & First Heart Field
Unclassified
Embryo Age:
E7.75
E8.25
E9.25
UMAP 1
UMAP 2
B
Mab21l2
Tbx5
Tbx1
Osr1
2.0
1.5
1.0
0.5
0.0
C
pseudotime
E
D
F
Aplnr
Cfc1
Tmem88
Isl1
Mab21l2
Tbx5
Fgf10
Tbx1
Mef2c
Nkx2-5
Nr2f2
Myh6
Myh7
Myl2
Shox2
Tbx18
Normalized Expression
Pseudotime

**Supplementary Figure 3. Representation of the in vivo mouse embryo heart scRNA-seq dataset with different variables**

A. UMAP representation of the mouse embryo heart scRNA-seq dataset with superimposed colours representing cell lineages (Left) and embryo age (Right).
B. Single cell expression of JCF (*Mab21l2*), FHF (*Tbx5*), SHF (Anterior: *Tbx1*, posterior: *Osr1*) used to segregate the two cardiac lineages in the dataset.
C. UMAP representation of the JCF lineage superimposed with the pseudotime function and green dot representing the root node.
D. Dot plot illustrating individual expression of cardiac progenitor, heart field lineages, cardiomyocyte and their subtypes and epicardial markers across the JCF lineage pseudotime function.
E. UMAP representation of the SHF lineage superimposed with the pseudotime function and green dot representing the root node.
F. Dot plot illustrating individual expression of cardiac progenitor, heart field lineages, cardiomyocyte and their subtypes and epicardial markers across the SHF lineage pseudotime function.

| Gene | Forward Sequence | Reverse Sequence |
|---|---|---|
| *ACTN2* | GTACGTCTCTTGCTTCTACCAC | CTTCCATCAGCCTCTCATTCTC |
| *CFC1* | GCTGAAGCACTGGGTGAATA | GTTGTTCTGCTGTCTCTACCTAC |
| *COL1A2* | CCCAGCCAAGAACTGGTATAG | CCTTGGAAGTCACTCCTTCTAC |
| *COL3A1* | CTGGCATTCCTTCGACTTCT | AGCTTCAGGGCCTTCTTTAC |
| *FOXC2* | CCGACCCAACCAGACAATTA | GTTACCTGCGCTCTTCACA |
| *GAPDH* | CAAGAGCACAAGAGGAAGAGAG | CTACATGGCAACTGTGAGGAG |
| *HAND1* | CAAGGATGCACAGTCTGGCGAT | GCAGGAGGAAAACCTTCGTGCT |
| *HOXB2* | CTTCCCGACCTCAACTTCTTC | CACAGAGCGTACTGGTGAAA |
| *NKX2-5* | CACCTCAACAGCTCCCTGAC | AATGCAAAATCCAGGGGACT |

**Supplementary Table 1. RT-qPCR primer sequences utilized in this study.**
