## Supplementary material for "APLNR marks a cardiac progenitor derived with human induced pluripotent stem cells": Key Resources Table

| **REAGENT or RESOURCE** | **SOURCE** | **IDENTIFIER** |
| --- | --- | --- |
| Antibodies | | |
| Mouse monoclonal anti-APJ | R&D Systems | Cat#MAB8561 |
| Mouse monoclonal anti-cTnT | ThermoFisher Scientific | Cat#MA5-17192, RRID: AB_2538663 |
| Rabbit monoclonal anti-KDR | Cell Signalling Technology | Cat#9698S, RRID: AB_11178792 |
| Rabbit polyclonal anti-PDGFRA | Cell Signalling Technology | Cat#3164S, RRID: AB_2162351 |
| Rabbit polyclonal anti-COL3A1 | Novus Biologicals | Cat#NB600-594, RRID: AB_10001330 |
| Goat polyclonal anti-mouse IgG microbeads | Miltenyi Biotec | Cat#130-048-401, RRID: AB_244360 |
| Goat polyclonal anti-mouse IgG, IgM (H+L), Alexa Fluor 488 | ThermoFisher Scientific | Cat#A-10680, RRID: AB_2534062 |
| Goat polyclonal anti-rabbit IgG (H+L), Alexa Fluor 555 | ThermoFisher Scientific | Cat#A-21428, RRID: AB_141784 |
| Rat monoclonal anti-mouse IgG_1_, FITC | BioLegend | Cat#406605, RRID: AB_493292 |
| Goat polyclonal anti-rabbit IgG (H+L), Alexa Fluor 488 | Abcam | Cat#ab150077, RRID: AB_2630356 |
| Chemicals, peptides, and recombinant proteins |  |  |
| Geltrex LDEV-Free, hESC-Qualified, Reduced Growth Factor Basement Membrane Matrix | Gibco | Cat#A1413302 |
| StemFlex Medium | Gibco | Cat#A3349401 |
| Y-27632 Dihydrochloride | BioGems | Cat#1293823, CAS: 129830-38-2 |
| StemPro Accutase Cell Dissociation Reagent | Gibco | Cat#A1110501 |
| Matrigel Matrix LDEV-Free | Corning | Cat#356234 |
| StemPro-34 SFM | Gibco | Cat#10639011 |
| GlutaMAX Supplement | Gibco | Cat#35050061 |
| L-Ascorbic acid | Sigma-Aldrich | Cat#A4544, CAS: 50-81-7 |
| Human BMP-4 Recombinant Protein | Gibco | Cat#PHC9531 |
| Human Activin A Recombinant Protein | Gibco | Cat#PHC9561 |
| IWR-1-endo | STEMCELL Technologies | Cat#72564, CAS: 1127442-82-3 |
| TrypLE Express Enzyme (1X), no phenol red | Gibco | Cat#12604021 |
| Trypsin-EDTA (0.05%), phenol red | Gibco | Cat#25300062 |
| DMEM/F-12 | Gibco | Cat#11320033 |
| HBSS (10X), calcium, magnesium, no phenol red | Gibco | Cat#14065056 |
| Bovine serum albumin | Sigma-Aldrich | Cat#A9647, CAS: 9048-46-8 |
| RPMI 1640 Medium, GlutaMAX Supplement | Gibco | Cat#61870036 |
| B-27 Supplement (50X), serum free | Gibco | Cat#17504044 |
| TRIzol LS Reagent | Invitrogen | Cat#10296028 |
| iTaq Universal SYBR Green Supermix | Bio-Rad | Cat#1725121 |
| ProLong™ Gold Antifade Mountant | Invitrogen | Cat#P36930 |
| DAPI (4',6-Diamidino-2-Phenylindole, Dihydrochloride) | Invitrogen | Cat#D1306, CAS: 28718-90-3 |
| Critical commercial assays |  |  |
| BD Cytofix/Cytoperm Fixation/Permeablization Kit | BD Biosciences | Cat#554714 |
| QuantiTect Reverse Transcription Kit | QIAGEN | Cat#205311 |
| RNA 6000 Nano Kit | Agilent | Cat#5067-1511 |
| Chromium Single Cell 3’ Reagent Kit v3 | 10X Genomics | Cat# PN-1000092 |
| Deposited data |  |  |
| Single-cell RNA-sequencing | This paper | GSE174213 |
| Bulk RNA-sequencing | This paper | GSE174335 |
| Experimental models: Cell lines |  |  |
| Human male iPSC: hiPSC1 | This paper | N/A |
| Human female iPSC: hiPSC2 | This paper | N/A |
| Human female iPSC: hiPSC3 | This paper | N/A |
| OIigonucleotides | | |
| Primers for RT-qPCR, see Table S1 | This paper | N/A |
| Software and algorithms |  |  |
| Cutadapt | Martin, 2011 | https://github.com/marcelm/cutadapt/ |
| STARsolo | Kaminow et al., 2021 | https://github.com/alexdobin/STAR |
| Rsubread | Liao et al., 2019 | https://bioconductor.org/packages/release/bioc/html/Rsubread.html |
| scPred | Alquicira-Hernandez et al., 2019 | https://github.com/powellgenomicslab/scPred |
| Seurat | Butler et al., 2018 | https://github.com/satijalab/seurat |
| Monocle 3 | Bergen et al., 2020 | https://github.com/cole-trapnell-lab/monocle3 |
| topGO | Adrian Alexa, Jorg Rahnenfuhrer | https://bioconductor.org/packages/release/bioc/html/topGO.html |
| MAST | Finak et al., 2015 | https://www.bioconductor.org/packages/release/bioc/html/MAST.html |
| pySCENIC | Aibar et al., 2017 | https://github.com/aertslab/pySCENIC |
| FlowJo | N/A | https://www.flowjo.com/solutions/flowjo |
